# Bayesian detection of convergent rate changes of conserved noncoding elements on phylogenetic trees

**DOI:** 10.1101/260745

**Authors:** Zhirui Hu, Timothy B. Sackton, Scott V. Edwards, Jun S. Liu

**Affiliations:** Department of Statistics; Informatics and; Organismic and Evolutionary Biology, Harvard University, 1 Oxford Street, Cambridge, MA 02138, USA

## Abstract

Conservation of DNA sequence over evolutionary time is a strong indicator of function, and gain or loss of sequence conservation can be used to infer changes in function across a phylogeny. Changes in evolutionary rates on particular lineages in a phylogeny can indicate shared functional shifts, and thus can be used to detect genomic correlates of phenotypic convergence. However, existing methods do not allow easy detection of patterns of rate variation, which causes challenges for detecting convergent rate shifts or other complex evolutionary scenarios. Here we introduce PhyloAcc, a new Bayesian method to model substitution rate changes in conserved elements across a phylogeny. The method assumes several categories of substitution rate for each branch on the phylogenetic tree, estimates substitution rates per category, and detects changes of substitution rate as the posterior probability of a category switch. Simulations show that PhyloAcc can detect genomic regions with rate shifts in multiple target species better than previous methods and has a higher accuracy of reconstructing complex patterns of substitution rate changes than prevalent Bayesian relaxed clock models. We demonstrate the utility of PhyloAcc in two classic examples of convergent phenotypes: loss of flight in birds and the transition to marine life in mammals. In each case, our approach reveals numerous examples of conserved non-exonic elements with accelerations specific to the phenotypically convergent lineages. Our method is widely applicable to any set of conserved elements where multiple rate changes are expected on a phylogeny.

## Introduction

One of the major revelations of comparative genomics has been the discovery of regions of the genome falling well outside protein-coding genes that nonetheless exhibit considerable levels of conservation across evolutionary time (Bejerano et al. 2004; Siepel et al. 2005; Woolfe et al. 2005; Venkatesh et al. 2006; Lindblad-Toh et al. 2011). Changes in conservation of elements, such as conserved non-coding or non-exonic elements, in a subset of lineages is often associated with altered regulatory activity and ultimately phenotypic divergence (Mclean et al. 2011; Booker et al. 2016). Numerous studies have used changes in sequence conservation of conserved elements as means to identify regulatory regions which may be of particular importance for lineage-specific phenotypes. For example, Pollard et al. (2006) identified 202 regions accelerated in the human genome but conserved in other vertebrates, some of which are RNA genes and tissue-specific enhancers. Holloway et al. (2016) identified 4,797 regions accelerated at the base of therian mammals, many of which are noncoding and located close to developmental transcription factors. Booker et al. (2016) discovered 166 bat-accelerated regions overlapping with enhancers in developing mouse limbs, including one that likely regulated expression of the *HoxD* cluster important for forelimb development. Such studies demonstrate that noncoding elements play a crucial role in molding morphological diversity across diverse clades.

Phenotypic convergence, in which the same function or morphology evolves multiple times independently, often due to adaption to similar environmental changes, is generally believed to be a strong signature of natural selection (Kishida et al. 2007; Brawand et al. 2008; Stern, 2013; Meredith et al. 2014). However, we generally do not have a robust understanding of the genomic changes underlying phenotypic convergence (Wray, 2013; Rosenblum et al. 2014). Do convergent phenotypes arise from repeated use of the same underlying genetic elements, or do they arise via independent genetic pathways (Orr, 2005; Tenaillon et al. 2012; Parker et al. 2013; Storz, 2016)? Convergence at the molecular level can arise because of identical substitutions, and convergent rate changes can arise via consistent shifts in substitution rate in genomic regions influencing particular phenotypic targets of natural selection (Chikina et al. 2016; Partha et al. 2017). In this paper, we focus on detecting genomic regions with convergent shifts of substitution rate that are correlated with convergent phenotype changes. These regulatory elements are often quite short, ranging from a few base pairs to several thousand, and hence may require sensitive tools to detect branches of the tree with different substitution rates.

Statistical tests for rate changes along a phylogeny have been a part of phylogenetic methodology for several decades, and are closely tied to tests for a molecular clock (Huelsenbeck and Crandall 1997; Huelsenbeck and Rannala 1997). Several existing relaxed clock models (Drummond and Suchard, 2010; Heath et al. 2012) explicitly model substitution rate variation across phylogeny. The random local clock model implemented in BEAST (Drummond and Suchard, 2010) assumes an indicator variable for rate change at each node, enumerating all possible local clock configurations, and estimates the location and magnitude of rate changes between local clock regions on the phylogeny. These methods allow users to both estimate a phylogeny and divergence times while allowing for rate variation among lineages, but are less powerful at detecting evolutionary shifts in rate that are correlated with a specific phenotype change as they do not explicitly incorporate such correlations in the model. Moreover, although highly accurate and useful for validating various clock models, these methods are not easily scalable to genome-wide data such as is typically encountered when testing for rate changes in conserved elements across a clade for which whole-genomes have been sequenced (McLean et al. 2011; Booker et al. 2016; Holloway et al. 2016).

Molecular clock tests are closely linked to tests for associations between rate shifts and specific convergent phenotypes on the tree, and several recent methods have been proposed to identify these associations. The “Forward Genomics” method (Hiller et al. 2012; Prudent et al. 2016) tests the significance of the Pearson correlation between normalized substitutions and hypothetical phenotypic states on each branch. Chikina et al. (2016), studying protein-coding genes with convergent substitution rate shifts in marine mammals, quantified the difference of relative substitution rates between “terrestrial” and “marine” branches using a non-parametric test (Wilcoxon rank sum test); the same approach has subsequently been extended to non-coding regions (Partha et al. 2017). The PHAST method (Hubisz et al. 2011) tests a model allowing substitution rates shift in a specified subset of branches against a null model with constant rate for all branches using likelihood ratio. However, these methods for detecting genomic regions with parallel substitution rate changes are generally limited to testing a single pre-specified shift pattern on a phylogeny, which is less powerful since many possible shift patterns would be congruent with a correlation with phenotypes of extant species. Furthermore, as we show here, these methods do not always distinguish among strong acceleration in a single tip branch, weaker acceleration across multiple clades and acceleration on lineages other than the target lineages.

Other model-based methods do not require pre-specified shift patterns, but have other limitations. Coevol (Lartillot and Poujol, 2011) jointly models parameters of substitution process (e.g. substitution rates) and continuous phenotypes as a multivariate Brownian process and outputs the posterior distribution of the correlation between phenotypes and substitution parameters, but does not allow for discrete phenotypes. To our knowledge, the only model-based method for associating molecular rate changes with discrete phenotypes and which considers multiple patterns of rate/character transitions is TraitRate (Mayrose and Otto, 2011; Levy Karin et al. 2017). This method models the probability of rate shifts along a fixed ultrametric tree, an approach that was shown to deteriorate the performance in practice (Mayrose and Otto, 2011). Additionally, TraitRate only estimates the likelihood ratio indicating the association between sequence evolution and a given trait; it does not model the pattern of shifts in substitution rate explicitly.

Here, we introduce PhyloAcc, a Bayesian method to model multiple substitution rate changes on a phylogeny. PhyloAcc does not require pre-determination of the history of rate shifts but instead relies on estimating the conservation state of each branch for a given element based on sequences of extant species. The method allows each genomic region tested to have a different pattern of shifts of substitution rate. Using Markov Chain Monte Carlo (MCMC) (Liu, 2001) to sample from the posterior distribution, PhyloAcc outputs the most probable evolutionary pattern of rate shifts as well as its uncertainty for each genomic region. PhyloAcc also evaluates the strength of the association between rate shifts at a genomic region and phenotypic states using Bayes factors (Kass and Raftery, 1995). Unlike previous methods using maximum likelihood estimators of substitution rates and a single pattern of rate shifts, PhyloAcc considers the uncertainty of estimated substitution rates and all possible changes of substitution rates by marginalizing all nuisance parameters either numerically or analytically given the phenotypes of extant species. To demonstrate the power of PhyloAcc on real data, we apply the method to two classic examples of phenotypic convergence: loss of flight in birds (Mitchell et al. 2014; Sackton et al. 2018) and transition to marine life in mammals (McGowen et al. 2014; Foote et al. 2015; Chikina et al. 2016). In both cases, we use genome-wide data from hundreds of thousands of conserved elements to identify those elements with specific patterns of convergent rate shifts associated with our target phenotype, revealing novel, putative regulatory regions that may be repeatedly associated with these evolutionary transitions.

## New Approaches

### Hierarchical Bayesian Phylogenetic Model: Overview

The goal of our model is to identify branches on a phylogeny on which particular genomic elements change their substitution rate. We take as input a phylogenetic tree, with branch lengths representing the expected number of substitutions along each branch averaged across the genome. Such a starting tree is often available from phylogenomic studies and branch lengths can be estimated, for example, from a class of sites thought to be neutral, such as four-fold degenerate sites (Hubisz et al. 2011). We assume that the substitution process follows a standard continuous time Markov process. To model rate variation, we introduce the relative substitution rate per branch, *r*, such that the expected number of substitutions along a given branch for a given element will be *r* times the background average. Given many genomic elements of interest are relatively conserved and short in length, estimating substitution rates for each branch accurately is difficult, as the number of substitutions which are informative will often be very low. To overcome this challenge, we use a local clock model where the substitution rate of a given branch is expected to correlate with that of its parent branch, and hence will be informed by more substitutions.

We assume that, for each element, a limited number of rate categories occur on the phylogeny. We define ***Z***_*i*_ = (*Z*_*i*1_, *Z*_*i*2_, …, *Z*_*in*_) to denote the latent *conservation state* on each of *n* branches for element *i*; the substitution rate for each element, *r*_*i*_ on a branch depends on its latent conservation state. The transitions in **Z** are modeled as a Markov Chain, i.e. the state of a branch only depends on the state of its parent branch. The transition probability matrix of **Z** is denoted by *Φ*. Such a model permits independent gain and loss of conservation on multiple lineages and also encourages nearby branches to have the same state and substitution rate, which is reasonable for closely-related species and branches in a phylogeny and is also a common assumption in phylogenetics (Thorne, 1998; Rannala and Yang, 2007). The posterior distribution of **Z** indicates where changes of substitution rates occur in the tree, and the posterior ratio of substitution rates for each latent state indicates the magnitude of change. The procedure is illustrated in Figure 1 and Supplementary Figure S1.

**FIG. 1:**
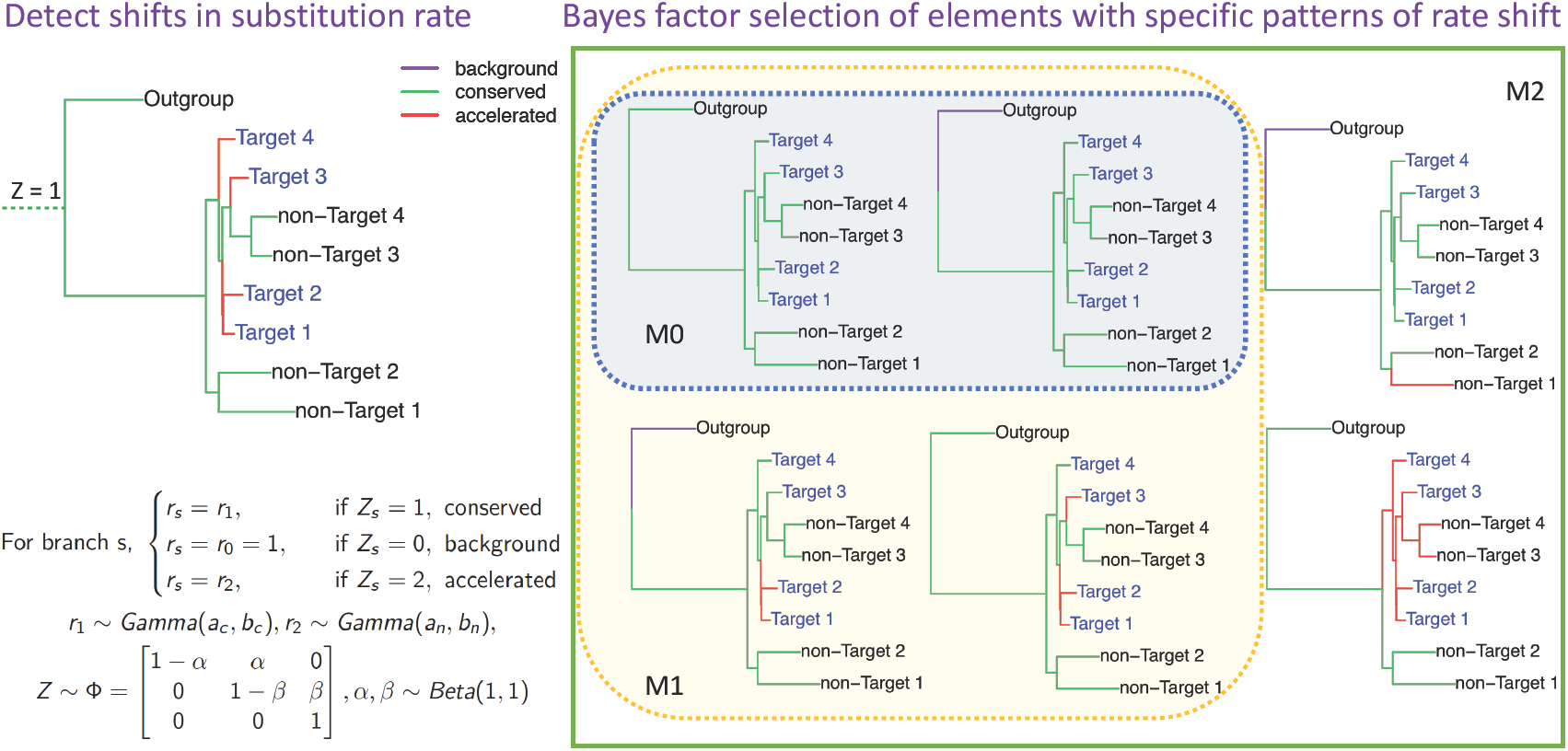
Illustration of the use of PhyloAcc to detect multiple accelerations and test hypotheses using Bayes Factors. The left panel shows the Bayesian phylogenetic model; right panel shows some examples of acceleration patterns in three nested models: null (M_0_), lineage-specific (M_1_) and full model (M_2_), respectively. Our method can recover shifts of substitution rate such as the top-left figure and select target-accelerated elements fitted by M_1_ (not M_0_). In the trees, target species are shown as blue; branch lengths represent the background substitution rates and branch colors indicate the latent states of substitution rate for a given element.

To test *a priori* patterns of substitution rate shifts that might be associated with phenotype changes, we compare the marginal likelihoods of three nested models by restricting the transition probability matrix *Φ* in some or all lineages: a null model without the specified shift pattern, a lineage-specific model satisfying the specific test pattern, and a full model allowing arbitrary shifts. We then compute two Bayes factors, 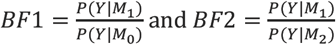, where *P*(*Y|M*_∗_) is the marginal likelihood of data set Y under model 0, 1 or 2, as criteria to identify DNA elements with specific evolutionary patterns. A larger Bayes factor implies stronger evidence from the sequence data in support of the specified pattern of rate shift (M_1_).

### Specific model for detecting multiple accelerations

We focus on a model of conserved element evolution, in which particular elements initially evolve at some background rate, become conserved at the root or some other branch on the phylogeny, and later potentially lose conservation in some lineages and thus evolve with an accelerated rate. We designate a conserved state with a lower substitution rate than sequences used to estimate background branch lengths (*r*_1_ < 1); an accelerated state with a substitution rate higher than that of the conserved state (*r*_2_ > *r*_1_); as well as a background state with the same substitution rate as those used in the input tree (*r*_0_ = 1, by definition). Informally, this model captures a process by which conservation arises as a transition from background to conserved state and is subsequently lost as when changing from the conserved to accelerated state, although the framework of PhyloAcc is flexible enough to model alternate scenarios. Note that in this model the accelerated rate is defined relative to the conserved rate, not the background rate, and thus accelerated rate can be less than, equal to, or greater than the defined background rate, as long as it is greater than the conserved rate. This allows us to model a variety of scenarios whose biological interpretation might include partially relaxed constraint or positive selection with a rate greater than background. Future extensions of phyloAcc with more rate categories could allow these alternatives to be distinguished. For example, it is straightforward to model multiple acceleration states in which a given element loses conservation deeper or more recent on a branch in independent clades, and thus have different numbers of substitutions realized on affected branches, or to allow for a loss of conservation state followed by a positive selection state with r >> 1. Although our model is flexible, it is ultimately agnostic as to the biological processes ascribed to the various estimated rate classes (see Discussion).

Formally, each branch is in either the background, conserved or accelerated state (i.e. Z_is_ ∈ {0,1,2}, *s* = 1,2, …, *n*), with substitution rates *r*_*i*0_ = 1, *r*_*i*1_ < 1 and *r*_*i*2_ > *r*_*i*1_, respectively for element *i* We will refer to branches in state Z_is_=2 with rate *r*_*i*2_ as “accelerated”, but we note that this acceleration is relative to the conserved rate, *r*_*i*1_. Such branches will always have rates greater than that of the conserved state, but could in principle have rates lower than the background rate *r*_0_ =1. Given that our candidate genomic regions (e.g. CNEEs) are mostly conserved, we expect that for any element most branches are in the conserved state. In practice, most branches in the accelerated state do indeed have estimated values of *r*_*i*2_> 1, with only a small percentage having values < 1.

To model how changes in latent conservation state occur along the phylogeny, we start by assuming that each element is in either the background or the conserved state at the root of the tree. In this way we can account for the fact that each element may not be conserved in all species in the tree, especially when distant outgroups are included. We assume that Dollo’s irreversible evolution hypothesis (Gould, 1970) holds for transitions from conserved to accelerated states, so that along each lineage *Z*_*is*_ can transit from a background to a conserved state, and then to an accelerated state but not the reverse (Felsenstein, 1973). By the irreversibility assumption, the transition probability matrix **Z**, has a simplified form: 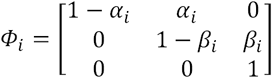, where *α*_*i*_ is the probability of gain conservation and *β*_*i*_ is the probability of loss of conservation. The model allows different transition probabilities *Φ* for each element, thereby allowing each element to be characterized by distinct evolutionary patterns.

As with many rate models, we assume a Gamma distribution for the priors of element-wise substitution rates (*r*_*i*1_ and *r*_*i*2_). Different hyperparameters of the priors for *r*_*i*1_ and *r*_*i*2_ distinguish conservation states. By adjusting the hyperparameters of the priors, the substitution rate for the accelerated state can be made stochastically higher than that of conserved state *a priori*. The hyperparameters of the priors are estimated by the sequence data from all elements in the data set, most of which will likely not change rate across the tree (Supplementary Text). This approach pools information from all elements to make estimates of substitution rates and latent states more reliable. Such pooling plays a larger role in cases where only a few branches are accelerated, and/or few substitutions occur per element. We used a Beta prior, which is conjugate to the likelihood function of **Z**, for the state transition probabilities (*α*_*i*_ and *β*_*i*_) for computational convenience.

In summary, the model has four parameters for each element: substitution rates in conserved and accelerated states (*r*_*i*1_ and *r*_*i*2_), transition probabilities to conserved and accelerated states (*α*_*i*_ and *β*_*i*_); and two latent random variables for each species or branch on the tree: the ancestral sequences and conservation states **Z.** Our method iteratively updates unobserved DNA sequences of ancestral species, latent states **Z**, substitution rates **r** and state transition probabilities *Φ* for each element by using collapsed Gibbs sampling (Liu, 1994) and adaptive Metropolis-Hasting algorithms (Roberts and Rosenthal, 2009) and outputs draws from the posterior distribution of **Z**. The resulting output can be used to reconstruct the sequence of shifts in rate, including the number of independent accelerations of a particular element, and quantifies the uncertainty of where in the tree (on which branch) accelerations occur. It also supplies posterior distributions of substitution rates **r**, indicating the magnitude of rate shifts (Supplementary Text).

### Testing parallel accelerations in target species

To test for an association between rate shifts and a set of pre-specified target lineages (phenotypically convergent species), for each element we compare marginal likelihoods between a null model assuming no acceleration in any lineage, and alternate models allowing either accelerations only in lineages associated with the convergent trait or accelerations in arbitrary lineages. In the null model (M_0_), all branches are in either the background or conserved state; in the lineage-specific model (M_1_), substitution rates on the branches leading to target species with the trait of interest can be accelerated while all other branches must be in either the background or conserved state; in the full model (M_2_), the latent conservation states **Z** can take any configuration across the phylogeny, in our implementation here, provided that Dollo’s irreversibility assumption on conservation states is not violated.

Formally, for M_0_ we restrict the transition probability from conserved to accelerated state (*β*) to be zero for all branches, and for M_1_ we restrict *β* = _0_ only to branches connecting the root to non-target species. In our applications, we assume *β* = _0_ for branches within outgroups in all three models such that all outgroup species cannot be in an accelerated state. To compare models, we compute the marginal likelihood *P*(*Y|M*_*i*_) for each model and compute two Bayes factors, BF1 and BF2 as described above, as criteria to identify DNA elements accelerated exclusively in target lineages. Including BF2 to identify elements with a specific evolutionary pattern is crucial to exclude elements accelerated in non-target species not associated with the specific phenotypic change, which might include regulatory elements with broader functions. The model and selection procedure are illustrated in Figure 1.

## Results

### Applications of PhyloAcc to examples of phenotypic convergence

To demonstrate the power of PhyloAcc, we conducted a simulation to compare the ability of PhyloAcc to distinguish various patterns of rate shifts where previous methods cannot. We then focus on two classic cases of convergent evolution: loss of flight in palaeognath birds (Mitchell et al. 2014; Sackton et al. 2018) and the transition to marine environments in mammals (McGowen et al. 2014; Foote et al. 2015; Chikina et al. 2016). We start by simulating data under the phylogenetic model for birds or mammals to verify the performance of our method, and then test for non-coding elements accelerated one or multiple times in flightless birds or marine mammals. We compared PhyloAcc with three alternative methods for selecting lineage-specific accelerated elements: phyloP in phast, which tests for clade-specific acceleration using a likelihood ratio test (Pollard et al. 2010), and two two-step procedures that first estimate branch-wise substitution rates using PAML (Yang, 2007) and then measure the correlation between rates and traits using either Wilcoxon rank sum test (denote as PAML+Wilcoxon) or phylogenetic ANOVA (Revell, 2012; denoted as PAML + phylANOVA).

### Simulation study: avian topology

To verify our ability to detect the correct evolutionary pattern, we simulated DNA elements with different patterns of rate shifts (i.e. different **Z**s) using a tree mirroring a recent phylogeny of birds (Jarvis et al. 2014), augmented by new genomes from palaeognathous birds (Sackton et al. 2018). Recent phylogenetic work supports the conclusion that the ratites (including ostrich, emu, cassowary, kiwi, rheas, and the extinct moas and elephant bird) are paraphyletic, implying convergent loss of flight in these lineages (Harshman et al. 2008; Baker et al. 2014; Mitchell et al. 2014; Yonezawa et al. 2017; Sackton et al. 2018). This scenario, in which target lineages are clustered in paraphyletic clade, is particularly challenging for existing methods. In most of our simulations, we set the length of each element to be 200 bp, which is about the median length in real data (Sackton et al. 2018). We also simulated one example with different element lengths to test the robustness of PhyloAcc. We generated 9 cases with different numbers of independent accelerations either within ratites and tinamous or neognath birds: 1) all branches are conserved; 2) only kiwi clade accelerated; 3) only ostrich accelerated; 4) only emu/cassowary branches accelerated; 5) only rhea clade accelerated; 6) all ratites accelerated except ostrich and moa; 7) all ratites accelerated; 8) both ratites and volant tinamous accelerated; 9) Five random species of non-ratite birds (Neognaths plus tinamous) accelerated (Supplementary Fig. S4). The total length of branches on which acceleration occurs increases from cases 1 to 8. We designed case 8 to demonstrate the specificity of our method, since the volant tinamou clade resides within the ratite clade, making it difficult to distinguish genomic elements accelerated from the ancestor of both tinamous and ratites from those only accelerated in ratites. In each case, we simulated 500 elements whose conserved and accelerated rates are generated randomly from gamma distributions (Supplementary Fig. S5C, Materials and Methods). The average proportion of nucleotides differences per element between pairs of species varies from 0.06 (case 1) to 0.13 (case 8). We first filtered out elements with *BF*2 < 1, a result indicating that species other than target lineages (ratites in this case) might be accelerated. We then ranked all other elements based on BF1. Similarly, each element was ranked and selected based on the test statistic or p-values output by other methods.

To test the sensitivity and specificity of our method in discerning target-specific accelerated elements from non-accelerated elements, we mixed the simulated accelerated elements with some non-accelerated elements from case 1 for each of the six ratite accelerated cases from cases 2–7. Receiver operating characteristic (ROC) curves resulting from varying selection thresholds are shown in Figure 2A. Not surprisingly, among different acceleration cases, all methods achieve a higher sensitivity when more accelerated target lineages are present. PhyloAcc consistently performed the best (except for the ostrich-accelerated case (case 3), in which phyloP is the best) at detecting elements accelerated among ratites based on BF1, whose distribution for different ratite-specific accelerated and background elements is shown in Supplementary Figure S6A. Since phyloP performs similarly or better than PAML+Wilcoxon and PAML + phylANOVA, we mainly focus on comparisons with phyloP, over a variety of sequence lengths: 100, 200, and 400bp (Fig. 2B). Both methods perform better with longer sequences and PhyloAcc is better for all lengths in most cases. PhyloAcc is also robust to different choices of the priors for substitution rates (Supplementary Fig. S7). Thus, under a variety of evolutionary scenarios, PhyloAcc has high power to detect lineage-specific rate shifts in conserved elements.

**FIG. 2:**
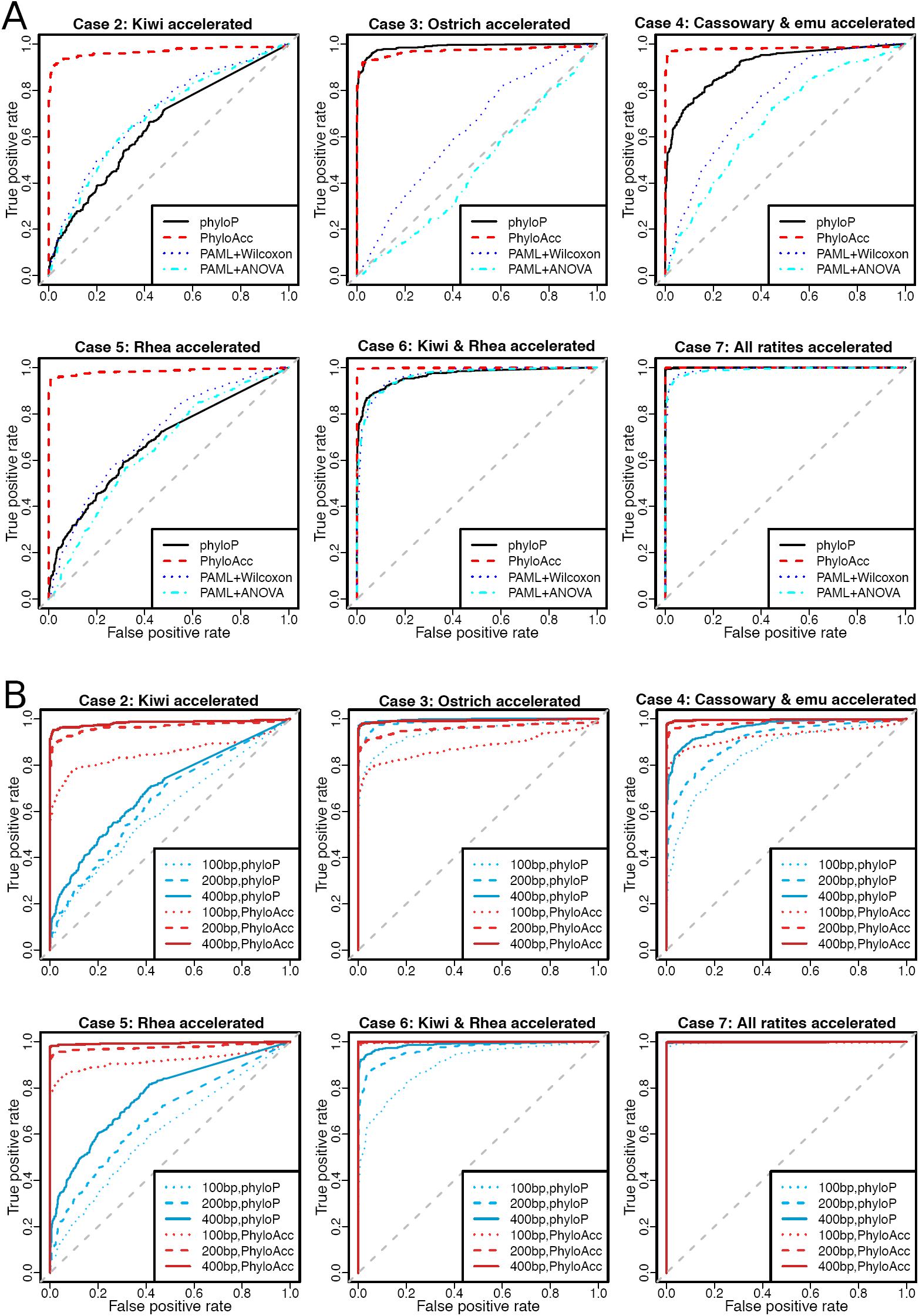
Simulated results on avian topology. (A) ROC curves for PhyloAcc, phyloP and PAML+Wilcoxon, PAML+phylANOVA in different ratite acceleration cases. (B) ROC curves for PhyloAcc and phyloP in different ratite acceleration cases and different lengths of elements. We treated elements with each acceleration pattern (case 2–7 separately) as positive and all conserved elements (case 1) as negative, and compared sensitivity and specificity of PhyloAcc to others.

Since phyloP can only test one pre-defined shift along a tree, a shift that moreover is usually not known with certainty *a priori*, we tested two common choices of accelerated branches: all tips of target species and all subtrees within target lineages, which is the acceleration pattern inferred by parsimony (Materials and Methods). With the second choice of accelerated branches, phyloP and PhyloAcc have comparable performances and phyloP is only better for shorter sequence in some cases (Supplementary Table S1).

To compare false positive rates (FPRs) of different methods, we mixed 100 elements from cases 2-9 together and with 5000 elements without acceleration from case 1, a situation that imitates the small proportion of target-specific accelerated elements in real data. Positive outcomes include cases 2-7, in which accelerations occur only within target species; other cases involve negative cases, in which either no acceleration occurs or accelerations occur outside target lineages. At 5% FPR, all methods except PhyloAcc could only identify a small number of true target-specific accelerated elements, because they incorrectly selected elements from case 8 as target-specific accelerated. In contrast, PhyloAcc successfully identified almost all the target-specific accelerated elements across all cases at low false positive rate (Supplementary Table S2).

The main reason for the superior performance of PhyloAcc in terms of controlling FPR is that PhyloAcc will not select elements accelerated in non-target lineages, e.g. in case 8 where acceleration occurs in the ancestors of ratites and tinamous. Other methods are not designed to control for this case; indeed the test statistic from phyloP or the two-step methods can be even larger in case 8 than in some target-specific accelerated cases. As an example, we showed the distribution of log-likelihood ratio output by phyloP compared with log-BF1 and log-BF2 by PhyloAcc in different cases (Supplementary Fig. S5B). In case 8, log-BF2 is less than -5 for 95% elements, because only the full model (M_2_), which allows for rate shifts on arbitrary branches, fits the data adequately. Thus, almost all elements are reported as not target-specific. By contrast, very few elements have log-BF2 less than 0 in cases 1-7, since the Bayes factors favor the simpler model if both models fit the data equally well (Supplementary Fig. S6B). Thus, our method achieves high specificity using BF2 as a filtering criterion. In additional simulations (Supplementary Fig. S5A), we show that our method has a lower false positive rate and a higher power in identifying elements with a shift in substitution rate within a set of species, and is thus well suited to identify either shared or independent rate changes.

### Inferring the pattern of acceleration of individual genomic elements

We confirmed that PhyloAcc can recover the true pattern of acceleration (pattern of latent states) for individual genomic elements by comparing the model estimated latent states to the true simulated values. For each simulated element, we compared the posterior probability of **Z** under the full model output from PhyloAcc with the true simulated pattern and defined the result as “correct” if the posterior probabilities of the true latent state on each branch are all above 0.7. Accuracy is then defined as the proportion of correctly detected elements. In our simulations, the ratio between accelerated and conserved rates is typically around 5 ∼ 1_0_, and the accuracy is above 60% in all cases. The accuracy is limited primarily by the posterior uncertainty of conservation state on short branches due to the lack of sufficient signal on those branches.

Previous methods selecting accelerated elements on particular branches do not always distinguish different patterns of acceleration from the score they produce. To illustrate this, we compared the log-likelihood ratio using phyloP and the p-value from Wilcoxon rank sum test (Pollard et al. 2010; Chikina et al. 2016) in the simulation cases above. Since these scores depend on both the magnitude and pattern of acceleration, it is not hard to find elements with the same log-likelihood ratios and p-values having either strong acceleration in a single lineage, multiple independent accelerations or a weak acceleration over an entire clade. As shown in Supplementary Figure S3, the second element is accelerated convergently in all target species (case 7) and thus more likely to be associated with the convergent phenotype. But the p-values of both likelihood ratio test and Wilcoxon rank sum test cannot distinguish between this pattern and one in which all ratites and tinamous are slightly accelerated (case 8) or in which only one lineage is accelerated (case 3). In contrast, PhyloAcc can identify both example 1 and 2 as ratite-accelerated elements, indicted by large log-BF1, but can also exclude element 3 because it has a negative log-BF2.

To compare the ability of PhyloAcc and other software to apply rate shifts to specific branches, we turned to BEAST2 (Bouckaert et al. 2014), since phyloP cannot output the pattern of acceleration on each branch. We designated the pattern detected by BEAST2 as correct if the posterior probability of rate shifts on branches with true state transitions is above 0.7 and below 0.3 for others. As seen in Figure 3, the accuracy of both methods increases as the ratio between accelerated and conserved rates increases, since the conservation state of short internal branches is easier to determine when we observe more substitutions, which will tend to occur when accelerated rates are high. In accordance with accuracy, for PhyloAcc, BF1 is also increases with the rate ratio, and BF2 stays below zero and decreases as *r*_2_/*r*_1_ grows in cases 8 and 9 (Supplementary Fig. S6C).

**FIG. 3:**
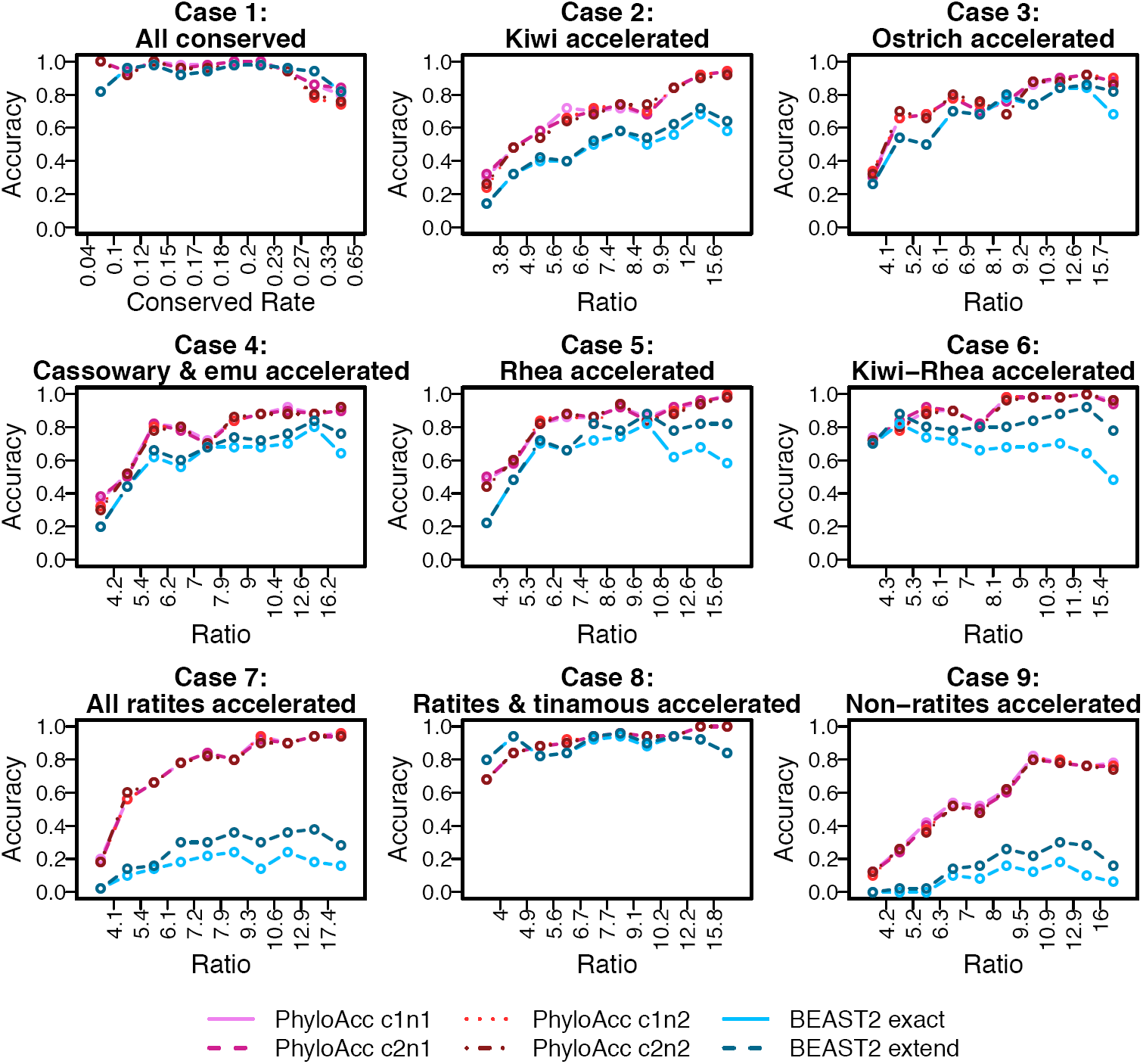
Comparison of accuracy recovering substitution rate shift patterns between BEAST2 and PhyloAcc in each simulation case. In each case, we ordered and categorized the simulated elements into 10 equal-sized groups according to the ratio between substation rates of accelerated and conserved states (the quantiles of *r*2/*r*_1_ in each group are shown in Supplementary Fig. S8A). X-axis shows the boundary of the ratio in each group; red curves are the accuracy of PhyloAcc (using different priors on substitution rates) and blue curves are of BEAST2. c1 and c2 are Gamma(5,0.04) and Gamma(1,0.2) respectively, narrow and wide prior for conserved rate; n1 and n2 are Gamma(10,0.2) and Gamma(4,0.5) respectively, narrow and wide prior for accelerated rate. “cXnX” means a combination of them. “BEAST2 exact” shows the accuracy recovering the true pattern; while “BEAST2 extend” shows the accuracy allowing “loss-regain” pattern.

BEAST2 has a comparable accuracy when no clade or a large clade is accelerated, but performs worse than PhyloAcc in cases with multiple independent rate shifts (e.g. cases 7 and 9) or with rate shifts on short branches (e.g., case 2). The model implemented in BEAST2 allows transitions between conserved and accelerated rates in both directions. As a consequence, BEAST2 tends to misplace the origin of an acceleration at a node deeper than the true node and then infer a regain of conservation in the clade whose rate is unchanged. For example, in the case in which only the rhea clade is accelerated (case 5), some elements are estimated as accelerated at the ancestor of rheas, kiwis, emu and cassowary, and then regain conservation in this clade except for rheas. There might be too few substitutions on these short internal branches for BEAST2 to determine their conservation state. Even when this type of “loss-regain” pattern inferred by the algorithm are counted as correct, PhyloAcc still performed better (Fig. 3). We also show that our model can recover the true conservation state with a high certainty (posterior of true latent state is around 1) for all but the shortest branches (Supplementary Fig. S8B and Fig. S9), and that it also appears robust to the presence of indels in the alignment (Supplementary Fig. S10).

### Simulation study: mammalian phylogeny

We next sought to validate our method in a second simulation study, this time focusing on the common scenario where a convergent phenotype arises in multiple, distantly separately lineages on a phylogeny. We used a recent tree for 62 mammals (Murphy et al. 2004), focusing on the transition to marine habit (Foote et al. 2015; Chikina et al. 2016), and simulated DNA elements under different patterns of substitution rates variation. We compared PhyloAcc with phyloP, PAML+Wilcoxon and PAML+phylANOVA in various cases: 1) all lineages conserved; 2) cetaceans (dolphin and killer whales) accelerated; 3) pinnipeds (seal and walrus) accelerated; 4) manatee, seal and dolphin accelerated, i.e. one species from each of the three independent lineages; 5) all five marine mammals accelerated; 6) pinnipeds and panda (sister lineage of pinnipeds) accelerated; 7) species descending from the common ancestor of cat and pinnipeds (Supplementary Fig. S11). Cases 2 through 5 are marine mammal-specific accelerated cases, whereas 6 and 7 is a case of non-specific acceleration. The average proportion of per-element nucleotides differences between pairs of species ranges from 0.09 (case 1) to 0.12 (case 7).

We conducted an analysis similar to that of the avian data set to compare the sensitivity and specificity of PhyloAcc and other methods for identifying substitution rate shifts. PhyloAcc has higher sensitivity to detect genomic elements accelerated within marine mammals than other methods, will exclude elements accelerated in species other than marine mammals by the BF2 criterion (Supplementary Fig. S12), and the FPR drops below 5% when selecting elements with log-BF2>0 and log-BF1>0 (Supplementary Fig. S12B). In combination with the avian simulation, these results suggest that the sensitivity and specificity of PhlyoAcc is expected to be high for a wide range of evolutionary scenarios regardless of the topology of target species on the phylogenetic tree.

### Detecting accelerated CNEEs in real data: avian case

We next applied PhyloAcc to detect ratite-accelerated conserved non-coding regions based on a set of 284,001 CNEEs identified in birds (Sackton et al. 2018). Using PhyloAcc, we identified 786 CNEEs with strong evidence for ratite-specific acceleration (log-BF1 > 20 and log-BF2 > 0), among which 80% have posterior median of accelerated rate greater than 1 under full model. The rhea clade is the most likely lineage to be accelerated among the 786 ratite-specific accelerated CNEEs, followed by kiwis, with the ostrich branch less likely accelerated among all ratites (Fig. 4A). PhyloAcc outputs the posterior probability of the conservation state on each branch, which are used to infer how many species of ratites are accelerated for each element as well as how many independent accelerations occurred within ratites (see Supplementary Text). Many of these CNEEs have experienced multiple independent accelerations within ratites: 54 (7%) CNEEs have three or more expected independent accelerations; 175 (22%) have been accelerated 2-3 times; and 480 (61%) have been lost 1-2 times (Supplementary Table S3; Sackton et al. 2018).

**FIG. 4:**
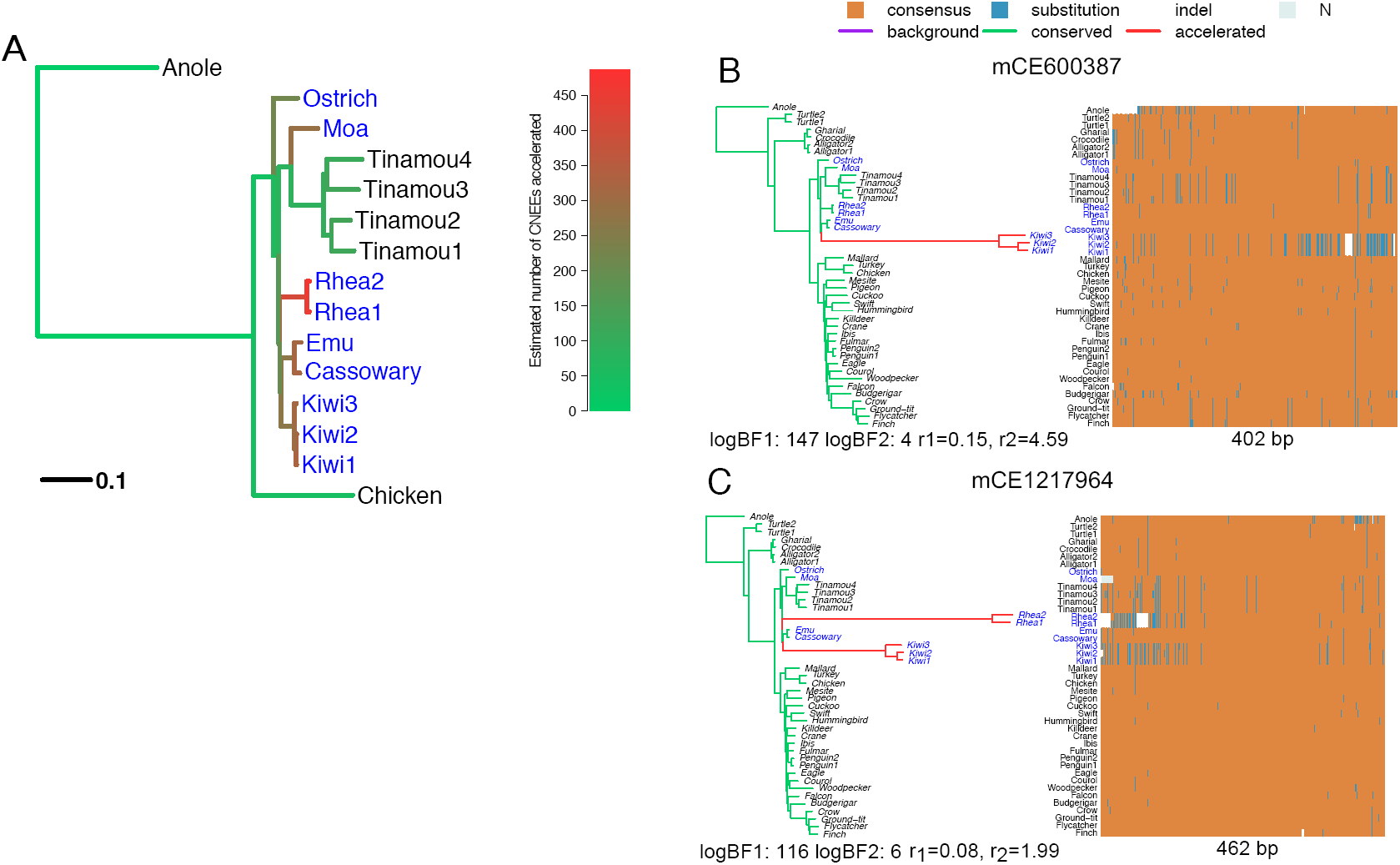
(A) Number of elements being accelerated per branch among ratite-specific accelerated CNEEs. Phylogeny for avian data set (only some species in neognathae and reptiles are shown for illustration). Palaeognaths consist of the flightless ratites and volant tinamous. Ratites are shown in blue. Branch lengths represent the background substitution rates. The gradient of the color indicates the expected number of elements being accelerated under the full model on that branch among 786 ratite-specific accelerated CNEEs. (B-C) Examples of ratite-accelerated CNEEs. For each element, the shift pattern of substitution rates under the full model is shown on the left represented by a phylogenetic tree with branch lengths proportional to the posterior mean of the substitution rate and colored by the posterior mean of **Z** (green is the conserved, red is the accelerated and purple is the background state). Longer and redder branch indicates acceleration occurred at a higher rate or earlier on the branch while shorter and greener one means later on the branch or no acceleration. Below the tree shows two log-BFs and conserved (*r*_1_)/accelerated rate (*r*2). In the sequence alignment heatmap on the right, each column is one position, each row is a species, and the element length is shown below. For each position, the majority nucleotide (T, C, G, A) among all species is labeled as “consensus” and colored as orange; others are labeled as “substitution” and colored as blue; unknown sequence is labeled as “N” and colored as gray; indels are shown as white space.

Among CNEEs that show strong signals of acceleration in ratites indicated by very large Bayes factors (BF1), many are accelerated in a single clade with one acceleration (e.g. mCE600387 accelerated only in kiwis, Fig. 4B) while some are accelerated in more than one clade (e.g. mCE1217964 accelerated in both rheas and kiwis, Fig. 4C). These are interesting candidate regulatory regions for further functional studies (e.g. Sackton et al. 2018).

### Detecting accelerated CNEEs in real data: mammalian case

As a second case study, we examined CNEEs accelerated in marine mammals. Though these mammals exhibit similar phenotypes upon transition to marine environments, the extent of molecular convergence in this system has been controversial, and largely focused on protein-coding genes (Foote et al. 2015; Chikina et al. 2016). Most genes with convergent sequence signatures are physiological and structural genes, with little evidence for convergent evolution in protein-coding genes controlling morphological adaptations, which may typically involve regulatory regions (Carroll, 2008). We applied PhyloAcc to 283,369 CNEEs identified from a whole genome alignment of 62 mammalian species, and identified 2106 elements showing evidence of substitution rate shifts specifically in marine mammals, with examples of acceleration on a single (Fig. 5A) and multiple clades (Fig. 5B). Compared with a random control group of mammals (aardvark, alpaca, camel, microbat, and David’s myotis bat) with no obvious shared characters but matching the topology of the five marine mammals (Supplementary Fig. S2B), we found more CNEEs showing substitution rate shifts in marine mammals than in control species (2106 for marine-accelerated vs. 1472 for control-accelerated elements with log-BF1>5 and log-BF2 > 5). We observed a larger Bayes factor between the lineage-specific model and the null model for marine-accelerated elements than for control-accelerated elements, indicating more dramatic changes of substitution rates affecting more species in marine mammals (Supplementary Fig. S14). In addition, more marine-accelerated CNEEs show parallel shifts in target lineages than controls: 696 (33%) of marine-accelerated elements show acceleration in 3 or more target lineages compared to 374 (25%) for control-accelerated elements (Supplementary Table S4); 93 (4.4%) of marine-accelerated elements show more than 2 independent accelerations compared to 33 (2.2%) for control-accelerated elements (Supplementary Table S5). To control for the chance that marine-accelerated elements are generally accelerated in more species, we compared the number of accelerated non-target species in each marine-accelerated CNEEs with that in control-accelerated CNEEs. We observed acceleration in only a small number of non-targeted species for marine-accelerated CNEEs and fewer than that in control-accelerated CNEEs (Supplementary Fig. S14).

**FIG. 5:**
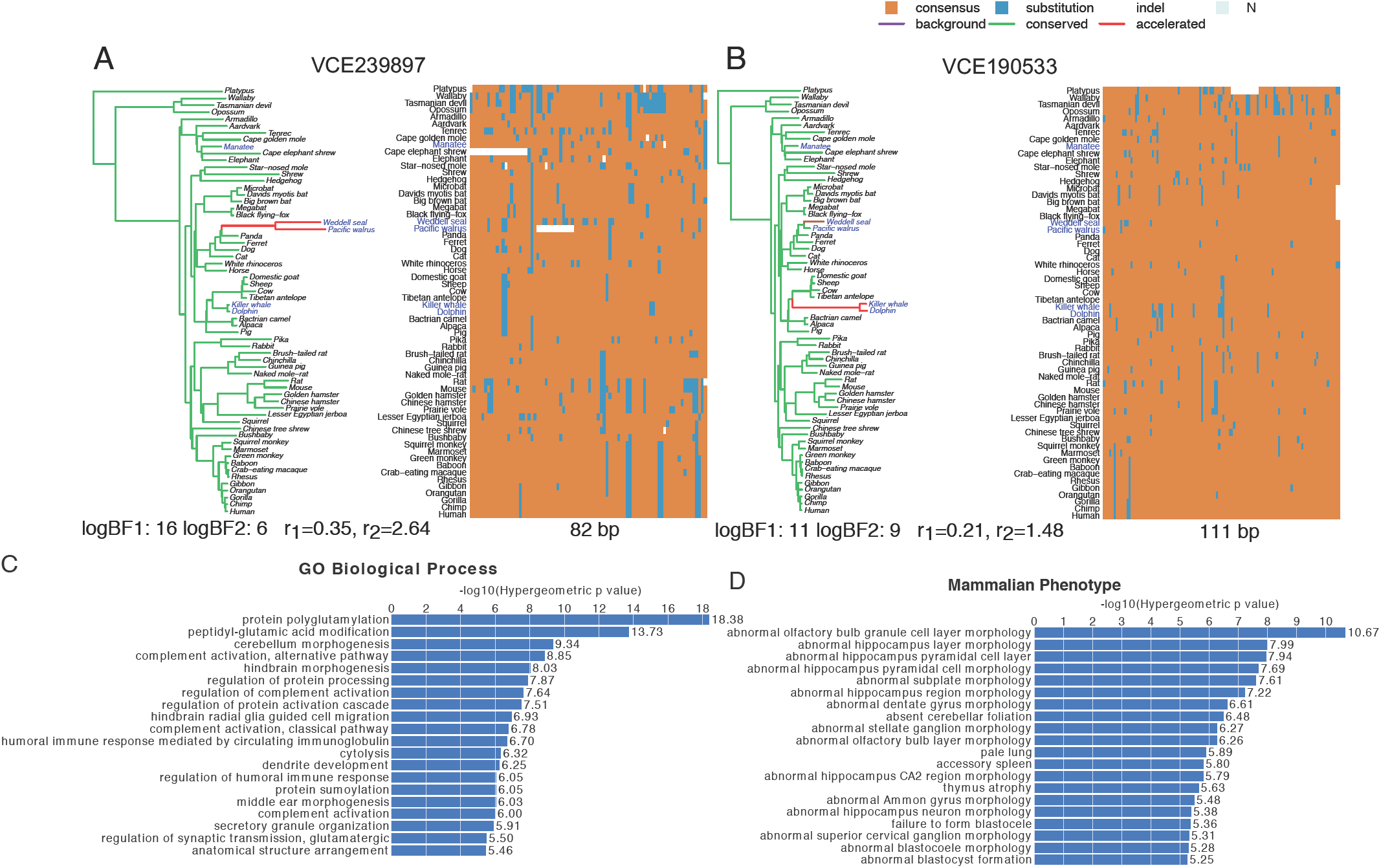
(A-B) Examples of marine mammal-accelerated CNEEs. For each element, the shift pattern of substitution rates under the full model is shown on the left represented by a phylogenetic tree with branch lengths proportional to the posterior mean of the substitution rate and colored by the posterior mean of **Z** (green is the conserved, red is the accelerated and purple is the background state). Longer and redder branch indicates acceleration occurred at a higher rate or earlier on the branch while shorter and greener one means later on the branch or no acceleration. Below the tree shows two log-BFs and conserved (*r*_1_)/accelerated rate (*r*_2_). In the sequence alignment heatmap on the right, each column is one position, each row is a species, and the element length is shown below. For each position, the majority nucleotide (T, C, G, A) among all species is labeled as “consensus” and colored as orange; others are labeled as “substitution” and colored as blue; unknown sequence is labeled as “N” and colored as gray; indels are shown as white space. (C-D) Enriched gene ontology (GO) terms (C) and mammalian phenotypes (D) of genes near marine-accelerated CNEEs. Only shown top 20 terms (all of them with FDR <0.01)

We tested for functional enrichment of genes near marine-accelerated CNEEs in mammalian genomes using GREAT (McLean et al. 2010). Among other functions, marine-accelerated CNEEs are predicted to regulate genes related to nervous and immune system including protein polyglutamylation, cerebellum morphogenesis, complement activation, and hindbrain morphogensis; these genes are also enriched in mammalian phenotype terms such as olfactory bulb granule cell layer morphology, hippocampus layer morphology, and subplate morphology (Figs. 5C-D). Many of the enriched functional terms are related to morphological traits, which reveals molecular adaptations overlooked by previous studies, which focused primarily on protein-coding genes. Checking individual genes associated with these enriched functional annotations, we found several genes surrounded by multiple top marine-accelerated CNEEs, including *TTLL3*, a beta-tublin polyglutamylase modifying microtubules and highly expressed in nervous system (Ikegami et al. 2006); *PROX1*, a member of the homeobox transcription factor family, associated with cerebellum morphogenesis; *C8B*, one component of the membrane attack complex, and in the complement pathway as part of the body1s immune response; *DAB1*, a key regulator of Reelin signaling pathway, playing an important role for neurogenesis; *KLF7*, a transcription factor, crucial for neuronal morphogenesis in olfactory and visual systems, the cerebral cortex, and the hippocampus (Laub et al. 2005); *FOXG1*, a transcription repressor, essential for brain development, especially for the region controlling sensory perception, learning and memory (Martynoga et al. 2005); and *GAS1* and *GLI2*, which function as transcription regulators in the hedgehog (Hh) pathway, important for embryogenesis (Martinelli and Fan, 2007). In contrast, control-accelerated CNEEs are enriched in only a few general gene ontology (GO) terms, such as cell fate determination, regulation of transcription and translation (Supplementary Fig. S15).

### PhyloAcc software

We implemented our method in the program PhyloAcc, which was written in C++, tested on Mac and Linux system, and is available at https://github.com/xyz111131/PhyloAcc. It requires as input: 1) a rooted phylogeny in .mod format (such as one produced by phyloFit in the PHAST package (Siepel and Haussler, 2004)); 2) a multiple alignment file concatenating sequences of all input (conserved) elements in FASTA format; 3) a bed file with the position of each individual element in the coordinate of concatenated alignment file (0-based); 4) and a parameter file. The .mod file should contain the transition rate matrix Q and the phylogenetic tree in Newick format with branch lengths (in units of substitutions per site) for background (neutral) sequences. The parameter file contains information on species names and parameters for the MCMC. For each element, PhyloAcc will output the posterior distribution of the latent conservation state (**Z**) for each branch, indicating neutral, conserved or accelerated states under the null, lineage-specific, and full models, respectively, and the marginal log-likelihood under each model as well as Bayes factors. The runtime of PhyloAcc compared to BEAST2 is shown in Supplementary Text. A detailed description of the usage as well as example simulation data sets and results are available in the GitHub repository. We also provide R scripts to generate figures summarizing the rate shift patterns as in this paper.

## Discussion

PhyloAcc provides a flexible framework to detect substitution rate changes along phylogenetic trees based on multiply aligned DNA sequences, conditional on annotated conserved sequence elements of interest (e.g. from PHAST or other tools). The method not only identifies DNA elements exhibiting changes of substitution rate in the lineages of interest, but also determines the branches, leading to either single or multiple lineages, experiencing changes of substitution rate (e.g. Sackton et al. 2018). We show here that PhyloAcc outperforms existing methods in simulations across a wide range of parameter space. Application to two biological datasets (loss of flight in ratites and shifts to marine habitat in mammals) revealed a number of noncoding elements accelerated independently on multiple phenotypically convergent lineages, suggesting that molecular convergence in regulatory regions may be commonly associated with phenotypic convergence.

The idea of matching sequence divergence profiles of either protein-coding genes or non-coding regions with repeated losses or gains of a given trait in multiple independent lineages to gain insight into the molecular basis of phenotype differences was first proposed as “Forward Genomics” by Hiller et al. (2012). Since then, this approach has been used in various groups of organisms, often yielding important insights into genome evolution and links between genotype and phenotype (Chikina et al. 2016; Prudent et al. 2016; Berger et al. 2017; Partha et al. 2017; Roscito et al. 2017). Compared to previous methods testing a pre-defined evolutionary history, our method can distinguish genomic elements with multiple independent accelerations within a target lineage from a single strong acceleration across the entire lineage. Our method also achieves a low false positive rate by contrasting the marginal likelihoods of models either allowing or prohibiting acceleration outside target species. Moreover, by averaging over the parameter space in competing models, Bayes factors offer a method of identifying accelerated elements that is more robust than previous two-step procedures and the likelihood ratio test implemented in phyloP. Two-step procedures that first use point estimates of branch-wise substitution rates and then test the correlation between rates and phenotypic traits often ignore the uncertainty in estimating branch-rates, which is quite uneven across different branches. The likelihood ratio test between constant rate and two-rates models is not appropriate in some extreme cases, such as when only a few branches have many substitutions.

The core utility of our software PhyloAcc is to detect changes of substitution rate on a tree for many conserved DNA elements, yielding the posterior distribution of substitution rates per conservation state (i.e. *r*_1_ and *r*_)_) for each element on each branch, from which the direction of rate change can be inferred. In the current implementation, we assume the same substitution rate for all accelerated branches, although our model can be extended to allow for different acceleration rates for each independently evolving clade. By introducing additional latent states, this extension can also allow for models distinguishing simple loss of conservation from acceleration due to natural selection or increased mutation rate. In addition, via Dollo’s assumption of irreversibility, our model allows at most two shifts on the tree for each lineage on the phylogeny, which may not be efficient for detecting elements that regain conservation after an ancient episode of adaptation. To relax this assumption, we could adjust the transition probability matrix of conservation states (**Z**) to allow for a small probability of transition from accelerated to conserved state. However, in many scenarios, the Dollo’s assumption is helpful, since the sequence data of extant species often do not provide enough information to distinguish consecutive substitution rate changes with opposite directions from no change at all, as illustrated in the simulation section when comparing with BEAST2. Additionally, the marginal likelihood is more difficult to compute for more complex models. Without doubt, our model is more specific in its goals than those implemented in more general phylogenetic packages (Drummond et al. 2006; Drummond and Suchard, 2010; Heath et al. 2012; Ogilve et al. 2017). As currently implemented, it is efficiently tailored to the one goal of detecting rate shifts in genome-wide noncoding elements on a phylogenetic tree. Further extensions of our framework would allow detection of rate shifts in protein coding regions, with advantages to distinguishing various closely related scenarios similar to those as studied here.

Although our method accounts for variation in background substitution rates across the tree, the background rate is likely not constant across different regions of the genome, a pattern that may impact our method and other previous methods (Hodgkinson and Eyre-Walker, 2011). However, we suspect that genome-wide variation in the local substitution rate is not a serious issue for our model, because our model detects acceleration relative to the conserved substitution rate at each genomic locus. Though the rate of background substitution is constant across the genome in our model, in our examples, only a few outgroup lineages are in typically in the background state, so the actual value of the background rate has relatively little impact on the estimated foreground rates. Nevertheless, the priors of substitution rate should reflect genome-wide variation in this background rate. For example, for a mutational cold-spot, both the conserved and accelerated rate might be low relative to the genome-wide average; if the prior for the accelerated rate is too high, the accelerated state might not be detectable; whereas for a mutational hot-spot, if the prior for the conserved rate is too low, the conserved state might be identified as background or accelerated. One way to tackle this genome-wide rate heterogeneity is to adjust the input background substitution rates and branch lengths on the phylogenetic tree for different segments of the genome, though this may introduce a degree of arbitrariness in the decision as to how to segment the genome.

Another major issue not addressed by our model is the possible heterogeneity in the topologies of gene trees across elements and across the genome. Heterogeneity in the topology of gene trees is expected to occur, especially during rapid radiations (Edwards 2009). Mis-specifying the phylogenetic tree can lead to mis-estimation of substitution rates on gene trees (Angelis and dos Reis, 2015; Hahn and Nakhleh 2016; Mendes and Hahn 2016). Bayesian multispecies coalescent models like *BEAST and starBEAST2 consider gene tree variations and can give more accurate estimate of per-species substitution rates (Heled and Drummond 2010; Ogilvie et al. 2017). It is possible to combine the multispecies coalescent model with our framework to test the correlation between rate shifts and phenotypes per genomic region accounting for phylogenetic uncertainty due to gene tree error or incomplete lineage sorting. This can be done by summing the likelihood over all probable gene trees given a species tree under the multispecies coalescent model, and then comparing the marginal likelihood integrating out the uncertainty of gene trees under different hypotheses.

Although in our examples we focus on loss of conservation accompanied by faster substitution rates, we do not attempt to distinguish among various processes — different types of mutation, increased mutation rates or natural selection — producing a specific pattern. GC-biased gene conversion (gBGC) is one of the factors that can increase local substitution rates across the genome and is often a confounding factor for the detection of adaptive evolution (Duret and Galtier, 2009; Kostka et al. 2012; Capra et al. 2013). Focusing on the avian example, we observed that ratite-accelerated elements have a higher GC content in ratites (Supplementary Fig. S13), which suggests a role for gBGC in acceleration. Although current version of PhyloAcc focuses on detecting acceleration but not distinguishing between positive selection and other processes, to demonstrate an approach for accounting for gBGC, we extended our method to jointly model gBGC and the effect of natural selection on substitution rates, using the framework that Kosta et al. (2012) applied to phyloP (Supplementary Text). We found that ∼30% ratite-accelerated elements also exhibited evidence for gBGC in accelerated lineages. We also provide an extended version of PhyloAcc online that can decouple the effect of gBGC from that of selection.

Identifying functions of regulatory regions is still a challenging task. Linking patterns of sequence evolution from diverse species with organism-level phenotypes has the potential to shed light upon regulatory functions of conserved non-coding regions. Such methodologies are still in their infancy. For example, Marcovitz et al. (2016) used parsimony to reconstruct traits and genome transitions and estimate their correlations, but in doing so were unable to account for the uncertainty of conservation pattern estimated from the sequencing data or the probability of a chance match between sequence evolutionary history and phenotype. Our method can be extended to provide the probability of a match between evolutionary profile of genetic elements with presence/absent patterns of hundreds of traits to predict phenotype-genotype pairs, an extension of the “Reverse Genomics” approach (Marcovitz et al. 2016). Additionally, our model could be extended to study functionally related genomic regions (e.g. using a Dirichlet process as prior for **Z**) based on similar patterns in sequences to discover novel functional groups of loci that may or may not influence known physiological and morphological traits. For example, a recent method, “CLIME” (Li et al. 2014; Li et al. 2018), uses a Bayesian approach similar in spirit to our model to cluster phylogenetic profiles of proteins to identify functionally related proteins. Jointly modeling a group of functionally related genomic regions in different species will provide more comprehensive insight of the evolutionary history and functional interaction of regulatory regions (Marcovitz et al. 2017).

## Materials and Methods

### Data sources for bird and mammal CNEEs

We obtained a whole genome alignment of 42 species (birds and non-avian reptiles) for ratite-accelerated region detection from Sackton et al. (2018; see this paper for full details on data collection). Conserved regions in the genome alignment were called by PhastCons using the Phast package (Siepel et al. 2005). A total of 284,001 CNEEs were extracted as DNA regions not overlapping any exons and at least 50 bp in length. Sequence from the extinct moa was subsequently added to CNEE alignments based on a pairwise moa-emu whole genome alignment (see Sackton et al. 2018 for details). For the mammalian dataset, we started with the UCSC 100-way vertebrate alignment (Blanchette et al. 2004; http://hgdownload.soe.ucsc.edu/goldenPath/hg38/multiz100way/), removed all non-mammalian sequences, and then extracted sequence for 383,185 CNEEs in a fashion similar to that for birds (conserved regions identified by PHAST, each at least 50 bp and not overlapping any exons). The list of species is in Supplementary Material. We filtered out CNEEs with poor alignment quality in 62 mammal species if the length of alignment gaps was longer than 80% of the whole alignment in more than 50 species, yielding 283,369 candidate CNEEs. For both phylogenies, we obtained branch lengths, parameters in the rate matrix of the nucleotide substitution model (GTR, General Time Reversible) and equilibrium nucleotide frequencies from phyloFit (Sackton et al. 2018 or UCSC, respectively) using background, putatively neutral sequences (in our case fourfold degenerate sites; Siepel et al. 2005).

### Simulating DNA Sequences

We simulated DNA sequences according to the joint model of sequences and conservation states (equation (2) in Supplementary Text) using the same phylogenetic tree and estimated rate matrix Q from sequence alignments as in the avian or mammalian data set using an in-house program. For the ratite simulation, we simulated 500 elements in cases 2-9 and 5000 elements in case 1 with length 200-, 100- and 400-bp under different configurations of **Z.** For the mammal simulation, we simulated 500 elements (200bp each) for each case. The conserved rate *r*_1_ was sampled from *Gamma*(5,0.04) and the accelerated rate *r*_2_ was sampled from *Gamma*(15,0.1), which are about the range of conserved and accelerated rates observed from real data (Supplementary Fig. S5C and Fig. S12D).

### Detecting Substitution Rate Shifts using BEAST2

We used BEAST version 2.4 in our simulation. The control file was generated by Beauti v2.4.7 with a fixed tree topology and branch lengths, the parameters of the substitution model taken at the true values, and no rate variation across sites within an element (Supplementary material). To get the substitution rate shift pattern, we used the random clock model, which has a binary variable indicating changes of rate and a clock rate for each branch. The priors (Gamma distribution for clock rate and Poisson distribution for shifts) and MCMC updates (total 10^6^ iterations and saved every 10^3^ steps) are set as default. Finally, we discarded first 20% of steps as burn-in and extracted the posterior distribution of the indicator variable from the MCMC log file.

### Detecting Substitution Rate Shifts Using PhyloP

To compare the performance of different methods for selecting elements with specific acceleration patterns, we used the phyloP program from PHAST v1.3 with the options --features (bed file) --method LRT --mode ACC --branch (target species) in our simulations, which used the likelihood ratio test to detect acceleration in all ratites or marine mammals. For simulations on the mammalian phylogeny, we only specified the tip branches of marine mammals as accelerated, whereas for the ratite analysis we specified both tip branches and the clade containing all ratites, which requires the minimum number of accelerations (see Supplementary Text).

### Detecting Substitution Rate Shifts Using PAML

We first estimated the branch lengths for each element using baseml program in PAML (version 4.8). We used “no molecular clock model” option, GTR (REV) nucleotide substitution model, and homogenous rate across sites within a region (by setting alpha = 0), which are the same settings as our simulations. For details, see the control file in the Supplementary Material. Then, we did a nonparametric test (Wilcoxon rank sum test) and phylogenetic ANOVA (using phylANOVA function in the R package phytools), comparing the substitution rates in ratites with other species. We obtained the p-values or test statistics from both tests in each simulated case.

### Function Prediction of CNEEs Using GREAT

To predict the regulatory functions of CNEEs in mammalian data set, we first extracted the genomic coordinates of these CNEEs using the human (hg19) genome as reference. To associate CNEEs with nearby genes, we used the “Basal plus extension” (up to 500Kb) option in GREAT. Then, we compared genes associated with marine- or control-accelerated CNEEs to genes near all CNEEs (background) and searched for any functional enrichment in GO biological processes and mammalian phenotypes from MGI. We only retained annotation terms that contain more than 5 genes in total, including at least 2 genes associated with accelerated CNEEs and at least 1.5-fold enrichment of tested CNEEs over all CNEEs.

## Acknowledgements

We thank Ziheng Yang and Huw Ogilvie for valuable comments on the manuscript, and Alison Cloutier, Phil Grayson, and Shaoyang Ning and other members in JSL’s lab for helpful discussion. SVE and TBS were supported by the National Science Foundation (DEB-1355343 to S.V.E. and EAR-1355292 to Julia Clarke), JSL and ZH were supported in part by the National Science Foundation (DMS-1613035) and National Institutes of Health (R01 GM122080). The computations in this paper were run on the Odyssey cluster supported by the FAS Division of Science, Research Computing Group at Harvard University.

## SUPPLEMENTAL MATERIALS

**Title: Supplementary_Material.pdf**

Supplementary Text, tables and figures. (pdf file)

**Mammal_species.txt**

The scientific name, common name and UCSC genome assembly of all species on the mammalian phylogeny. (txt file)

**Avian_species.txt**

The scientific name and common name of all species on the avian phylogeny. (txt file)

**Beast_config.xml**

The configuration file for running BEAST2.

**PAML_baseml.ctl**

The configuration file for running PAML.

